# Maternal high-fat diet induces sex-specific changes to glucocorticoid and inflammatory signaling in response to corticosterone and lipopolysaccharide challenge in adult rat offspring

**DOI:** 10.1101/780783

**Authors:** Sanoji Wijenayake, Mouly F. Rahman, Christine M.W. Lum, Wilfred C. De Vega, Aya Sasaki, Patrick O. McGowan

**Affiliations:** Center for Environmental Epigenetics and Development, Department of Biological Sciences, University of Toronto, Scarborough, 1265 Military Trail, Toronto, ON, Canada; Department of Cell and Systems Biology, University of Toronto, Toronto, ON, Canada; Department of Psychology, Department of Physiology, University of Toronto, Toronto, ON, Canada

**Author notes:** Correspondence to: Patrick O. McGowan, University of Toronto, Scarborough Campus, 1265 Military Trail, Toronto, Ontario, Canada. M1C1A4. Authors contributed equally to the manuscript (Sanoji Wijenayake, Mouly F. Rahman).

**Keywords:** Maternal high-fat diet, maternal obesity, offspring, neuroinflammation, glucocorticoid signaling, transcript response, corticosterone, lipopolysaccharide

## Abstract

**Background:** Acute elevations in endogenous corticosterone (CORT) with psychosocial stress or exogenous administration potentiate inflammatory gene expression. Maternal obesity as a result of high-fat diet (HFD) consumption has been linked to higher basal levels of neuroinflammation, including increased expression of pro-inflammatory genes in the amygdala. These findings suggest that exposure to maternal HFD may elicit pro-inflammatory responses in the presence of an immune stressor such as lipopolysaccharide (LPS), a component of gram-negative bacteria, as well as acute elevated CORT.

**Methods:** Rat offspring were exposed to maternal HFD or control diet (CHD) throughout pre and postnatal development until weaning, when all offspring were provided CHD until adulthood. In adulthood, offspring were ‘challenged’ with administration of exogenous CORT, to simulate an acute physiological stress, LPS, to induce an immune stress, or both. qPCR was used to measure transcript abundance of CORT receptors and downstream inflammatory genes in the amygdala, hippocampus and prefrontal cortex, brain regions that mediate neuroendocrine and behavioral responses to stress.

**Results:** HFD female offspring exhibited elevations in anti-inflammatory transcripts, whereas HFD male offspring responded with greater pro-inflammatory gene expression to simultaneous CORT and LPS administration.

**Conclusions:** These findings suggest that exposure to maternal HFD leads to sex-specific alterations that may alter inflammatory responses in the brain, possibly as an adaptive response to basal inflammation.

## INTRODUCTION

The hypothalamic-pituitary-adrenal (HPA) axis regulates circulating glucocorticoid (GC) levels at baseline and in conditions of stress. In the brain, GC binding to glucocorticoid receptors (GRs) in the amygdala, hippocampus (HPC), and prefrontal cortex (PFC) send feedback signals to the hypothalamus to mediate HPA axis activation or inhibition (1,2). GRs also mediate inflammatory signalling in the brain. GC-GR binding elicits the expression of anti-inflammatory genes including mitogen-activated protein kinase phosphatase 1 (MKP-1) and NFκB-inhibitor alpha (IκBα). MKP1 and IκBα inhibits nuclear translocation of nuclear factor kappa beta (NFκB) and thereby reduce NFκB-mediated transcription of pro-inflammatory cytokines, including IL-6 (3,4). While physiological levels of GCs such as corticosterone (CORT) suppress inflammation, acute increases in levels of CORT potentiate pro-inflammatory processes (3,5–8). For example, chronic unpredictable stress potentiates lipopolysaccharide (LPS)-induced NFκB activation and pro-inflammatory cytokine expression in the frontal cortex and HPC. Likewise, higher circulating CORT levels are associated with a potentiation of LPS-induced pro-inflammatory signalling in the brain (3). GR antagonism by RU-486 has been found to blunt the potentiating effect of CORT-mediated stress on LPS-induced pro-inflammation in the frontal cortex and HPC (3,4,9). These findings indicate a positive correlation between CORT-GR binding and pro-inflammation. However, it is unknown how chronic alterations in HPA axis activity may affect the combination of immune stress and acutely elevated CORT.

Several studies have linked developmental exposure to high levels of saturated fat through the maternal diet with altered glucocorticoid signalling and HPA axis activity in adult offspring. High levels of saturated fats in maternal obesogenic diets induce inflammation and stress in the mother that reaches developing offspring both *in utero* and during the lactation through milk (10,11). In humans, maternal high-fat diets (HFD) are linked to metabolic disorders as well as anxiety disorders in children (12–18). In rodent offspring born to obese dams fed HFD, there is altered expression of GR and downstream inflammatory genes in brain regions that regulate the HPA axis (19–25). For example, our lab found increases in pro-inflammatory gene expression of NFκB and IL6 in the amygdala in basal conditions, along with downregulated serum CORT (19).

In the current study, we predicted that immune stress in combination with CORT administration would lead to larger potentiation of pro-inflammatory effects in adult offspring exposed to maternal HFD compared to CHD offspring. To investigate this, female and male rats exposed to maternal control or HFD during the perinatal period were administered exogenous CORT to simulate psychological stress, LPS to induce immune stress, or simultaneous CORT and LPS challenge in adulthood. Transcript abundance of CORT receptors, including GR and downstream inflammatory pathway genes, was measured in the amygdala, HPC, and PFC. Female and male offspring were examined separately due to prominent sex differences in body weight, endocrine and behavioral responses to maternal HFD, CORT, and LPS exposures reported previously (19,22,26–28).

## MATERIALS & METHODS

### Animal Care

All experimental protocols were approved by the Local Animal Care Committee at the University of Toronto Scarborough and were in accordance with the guidelines of the Canadian Council on Animal Care. The adult rat offspring used for this study were untested littermates of the same cohort of animals in previous studies published by our group (19,20). For breeding, 7-week-old adult male and female Long Evans rats were purchased from Charles River Canada (St. Constant, QC), and housed with same-sex pairs, and maintained on a 12:12 h light-dark cycle (lights turned on from 7:00 am to 7:00 pm) with *ad libitum* access to food and water. Female breeders were placed on either a high-fat diet (HFD, 5.24 kcal/g, n=15) consisting of 60% fat, 20% protein, 20% carbohydrate (D12492; Research Diets, Inc. New Brunswick, NJ.), or a control house chow diet (CHD, 3.02 kcal/g, n=14) consisting of 13.5% fat, 28.5% protein, and 58% carbohydrate (5001; Purine Lab Diets. St. Louis, MO) four weeks prior to mating, throughout gestation, lactation, and until weaning. Litters were weighed weekly during cage changes and were otherwise left undisturbed until weaning (post-natal day (PND) 21), when they were all placed on a CHD diet and housed in same-sex pairs. At adulthood (PND90), body weights were measured prior to CORT and LPS injections (Figure 1).

**Figure 1.**
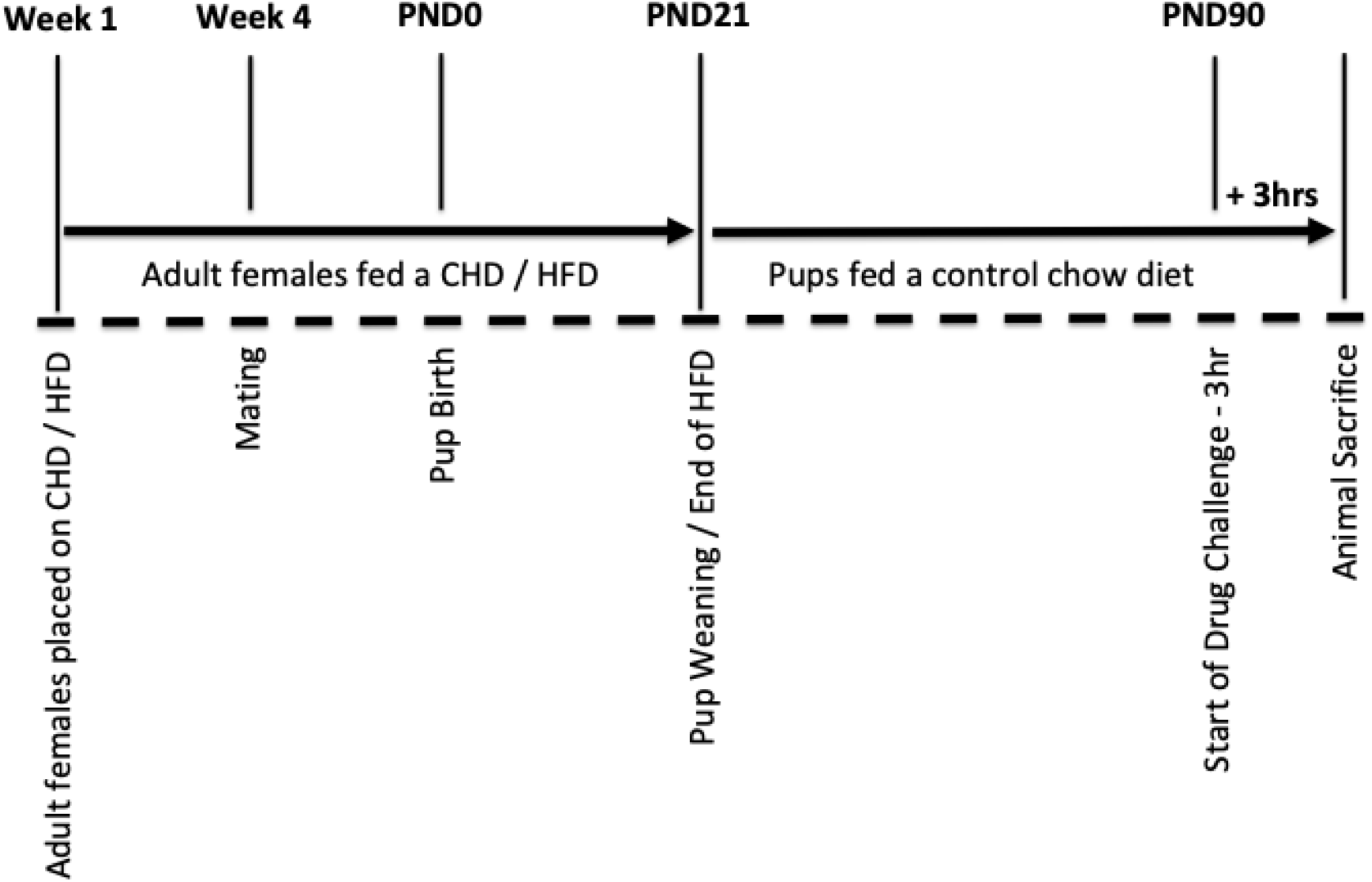
Overview of animal care and stress challenges. Adult female breeders were placed on either CHD (control house-chow diet) or HFD (high-fat diet) four weeks prior to mating and continued through gestation and lactation. Offspring were weaned on CHD at postnatal day (PND) 21. On post-natal day (PND) 90, offspring were injected with corticosterone (CORT), lipopolysaccharide (LPS), or CORT+LPS combination, and sacrificed 3 hours later.

### Adult CORT and LPS challenge

CORT dissolved in propylene glycol (27840; Sigma-Aldrich) and LPS from *Escherichia coli* O111:B4, (L2630; Sigma-Aldrich) were used for subcutaneous and intraperitoneal injections respectively. Adult female and male offspring were handled 2 min/day for five consecutive days starting from PND85. At PND90, animals were divided into one of four experimental groups; 1) subcutaneous dose of CORT (10 mg/kg of body weight), 2) an intraperitoneal dose of LPS (50 μg/kg of body weight), 3) a simultaneous dose of CORT and LPS (10 mg/kg, 50 μg/kg), or 4) handled controls (n=6 per diet, sex, and treatment) (29). A 10mg/kg dose of CORT was previously shown to lead to heightened anxiety-like behavior and circulating plasma CORT levels similar to that of several hours of acute physiological stress (30). The 50 μg/kg dose of LPS was shown to activate the HPA axis within 0.5-4 h, as shown by increased CORT levels in whole-blood (31) and induce pro-inflammatory cytokine expression in the hippocampus of offspring exposed to maternal HFD (22). Animals were sacrificed 3 h post injection by CO_2_ inhalation followed by rapid decapitation at the mid-point of the light phase (11-3 pm) to control for circadian-related changes in gene expression. Brains were dissected, flash-frozen in isopentane and dry ice, and stored at −80 °C.

### RNA Extraction and cDNA Synthesis

Whole amygdala, dorsal hippocampus, and medial prefrontal cortex were cryo-sectioned using a Leica CM3050 cryostat and stereotaxic coordinates (32). RNA was extracted from the amygdala, dorsal hippocampus, and medial prefrontal cortex using TRIzol reagent (15596026; Invitrogen) in combination with RNeasy Plus Mini Kit (74134; Qiagen) as per the manufacturer’s instructions. RNA quantification and quality assessments were done using a Nanodrop Spectrophotometer (ND-2000C; Thermo Scientific). 1 μg of total RNA was converted to cDNA using High Capacity cDNA Reverse Transcription Kit (4368814; Applied Biosystems) according to the manufacturer’s instructions.

### Gene expression Analysis by qPCR

Relative mRNA expression of glucocorticoid receptor (GR), mineralocorticoid receptor (MR), nuclear factor kappa light chain enhancer of activated B cells (NFκB), nuclear factor of kappa light polypeptide gene enhancer in B-cells inhibitor, alpha (IκBα), interleukin 6 (IL6), interleukin 10 (IL10), cluster of differentiation molecule 11B (CD11B), mitogen activated protein kinase phosphatase 1 (MKP1), and insulin-like growth factor 1 (IGF1) in the three brain regions was measured using a StepOne Plus real-time thermocycler with a Fast SYBR Green PCR master mix (4385612; Applied Biosystems). Primers were purchased from Qiagen or Eurofins Genomics and designed according to GenBank sequence information at the National Center for Biotechnology Information (NCBI) (Table 1).

**Table 1.**
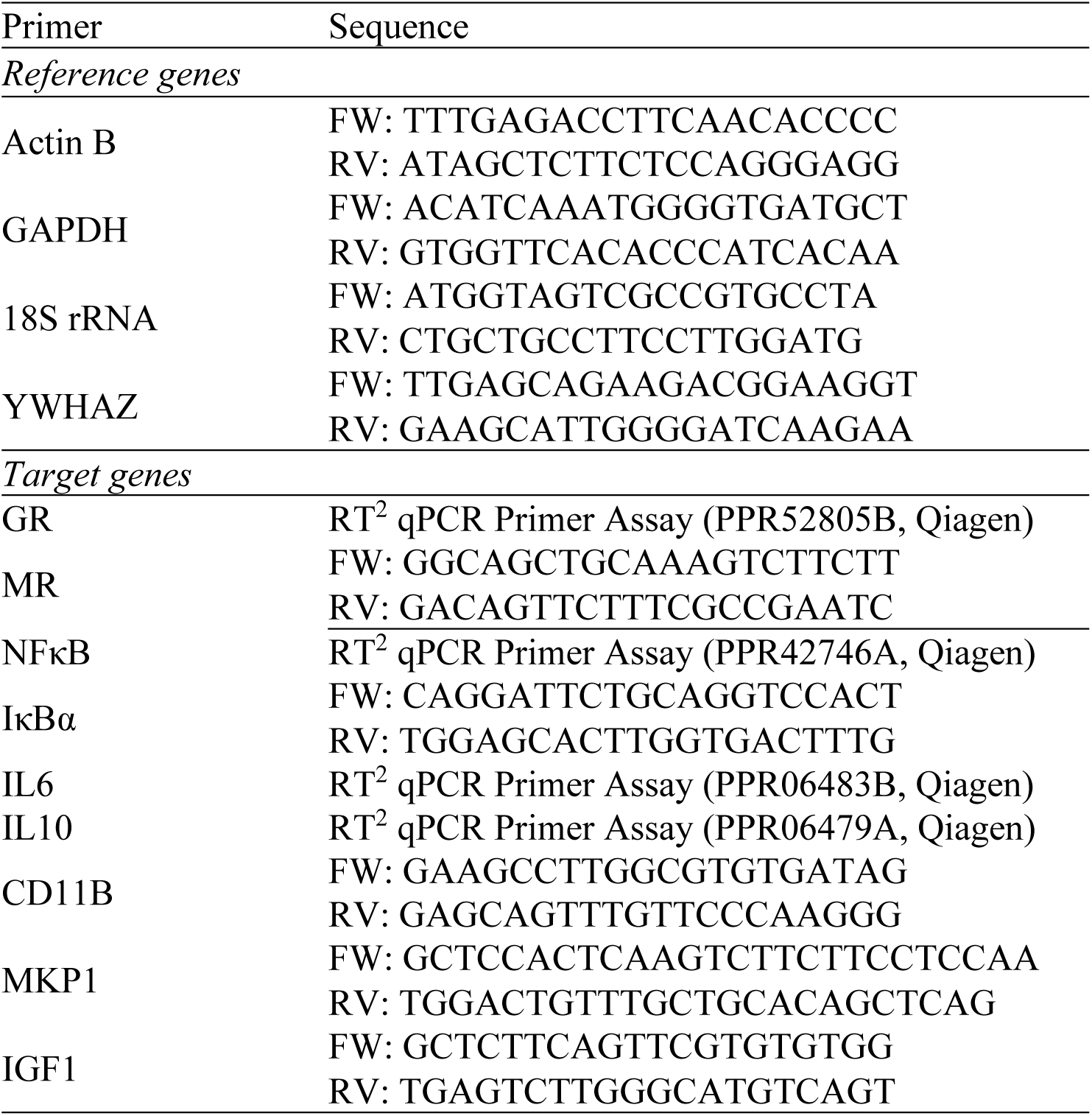
Primers used in qPCR analysis.

Relative gene expression was calculated using the quantity mean based on a standard curve of 11 serial dilutions ranging from 500 ng/μL to 0.49 ng/μL of cDNA. A standard curve was run per plate and per set of comparisons. Quantity means were normalized against the GEOmean of four reference genes, YWAZ, GAPDH, 18s, and Actin B. Reference genes were identified as stable internal controls based on geNORM analysis of stability across experimental groups, brain regions, and sex (33). Stability M values calculated by geNORM: YWAZ = 0.322, Actin B = 0.49, GAPDH = 0.49, 18S = 0.68. Relative transcript levels were expressed as mean ± SEM representing n=6 biological replicates per experimental condition.

### Statistical Analysis

Data analysis was carried out using SPSS (IBM Corp) and R Statistical Software (R Foundation for Statistical Computing, Vienna, Austria, 3.4.2). Adult offspring bodyweight was analyzed by 3-way (diet x drug x sex) ANOVA. qPCR data were analyzed within sex and brain region. A Shapiro-Wilk test was used to assess normality for all transcript data as the n<30. All transcript data were normally distributed and as such parametric analyses were carried out. Outliers were examined using boxplots and only extreme outliers with interquartile range of 3 or more were removed from the dataset. General linear model (GLM) univariate analysis was used to test for main effects of diet, challenge, and diet x challenge interactions. The Scheffe post-hoc test was used to conduct mean pairwise comparisons between diet and challenge groups. Relationships were considered statistically significant at p≤0.05.

## RESULTS

### Offspring body weight

As reported previously by our group for animals in this cohort, offspring with maternal HFD exposure showed no differences in body weight at birth, however weighed more than CHD offspring through weaning. In adulthood (PND90), male offspring weighed more than female offspring (main effect of sex (F_(1,93)_ = 220.20, p<0.001, Figure 2), and both sexes with maternal HFD exposure weighed more than CHD offspring (females (F_(1,46)_ = 19.71, p< 0.001, males (F(1,45) = 39.82, p< 0.01, Figure 2). There were no significant differences in body weight between animals assigned to any of the treatment groups (CORT, LPS, CORT+LPS, control handled).

**Figure 2.**
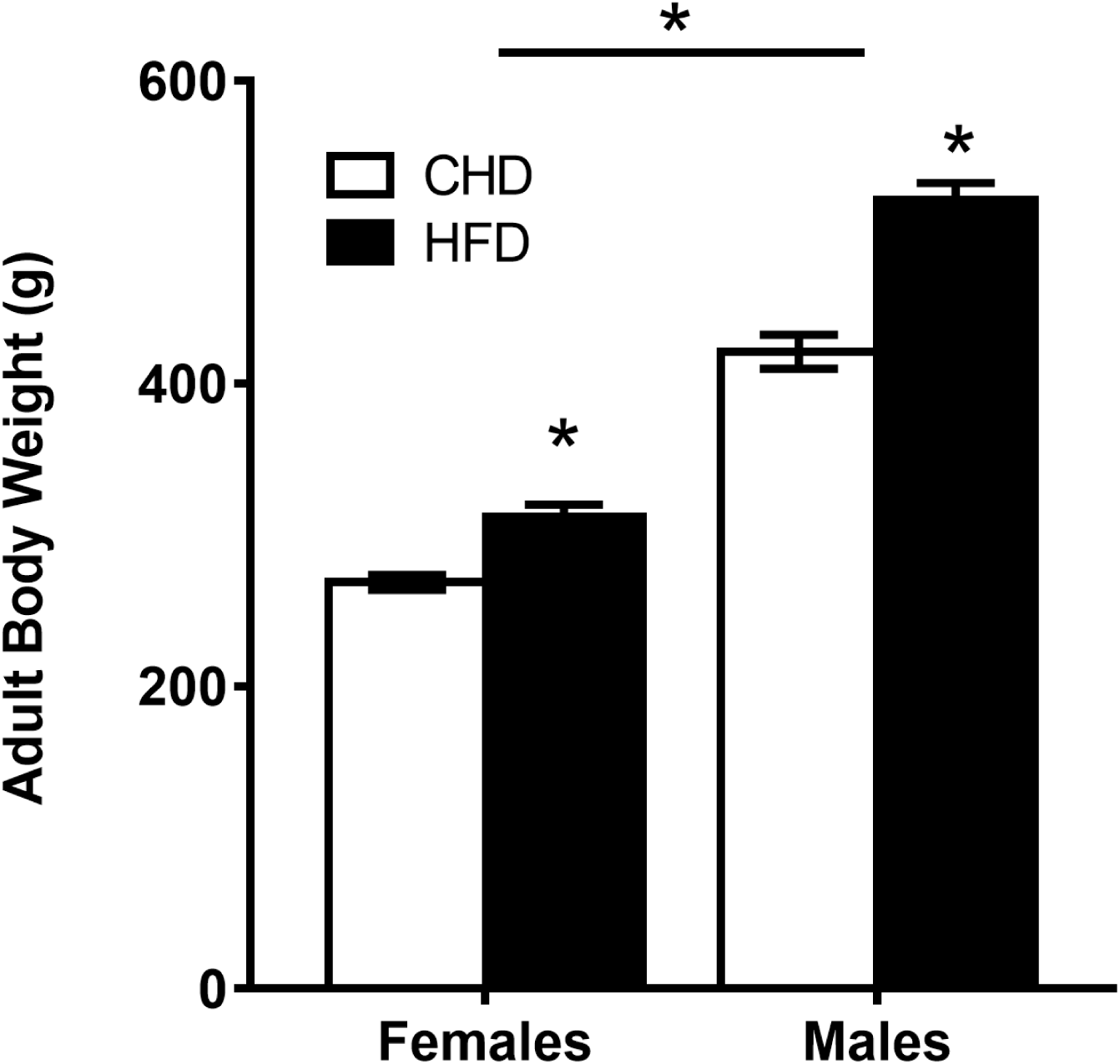
Offspring body weight in adulthood. Average body weight for PND90 offspring per sex and maternal diet condition ± SEM. n=24 for each group, CHD = control house chow diet, HFD = high-fat diet. *p < 0.0001 for main effect of diet, and main effect of sex.

### Transcript response to endocrine and immune challenge

#### CORT challenge

In female offspring, CORT challenge led to few differences in transcript abundance between HFD and CHD offspring (Figure 3A, 3B). In the amygdala, IκBα increased in HFD offspring (main effect of challenge (F_(3,19)_ = 9.63, p<0.01), Scheffe post-hoc p=0.015), but did not change among CHD offspring (Scheffe post-hoc p= 0.195, Figure 3C). In the hippocampus, IκBα increased in both diet groups (main effect of challenge (F_(3,20)_ = 13.35, p<0.01), Figure 3D). Also in the hippocampus, MKP1 increased in HFD offspring (main effect of challenge (F_(3,20)_ = 12.75, p<0.05), Scheffe post-hoc p=0.001), but remained unchanged in CHD offspring (Scheffe post-hoc p=0.985, Figure 3E).

**Figure 3.**
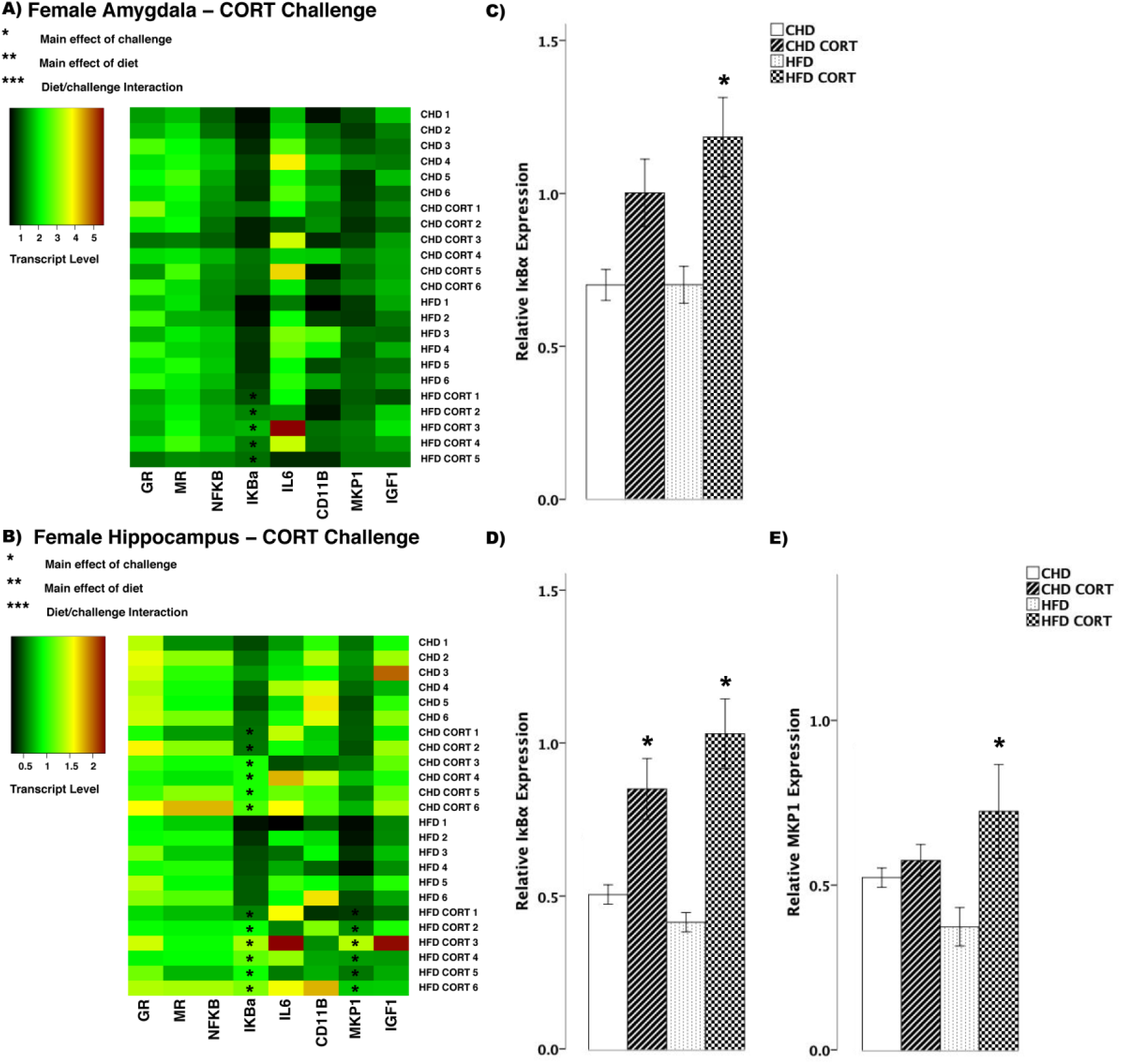
Transcript response to corticosterone (CORT) in the amygdala and hippocampus of adult females. **A and B:** Heatmaps represent transcript levels in the amygdala and hippocampus for individual animals. **C:** relative transcript abundance of IκBα (nuclear factor of kappa light polypeptide gene enhancer in B-cells inhibitor, alpha) in the amygdala. **D-E:** relative transcript abundance of IκBα, and MKP1 (mitogen activated protein kinase phosphatase in the hippocampus. Data presented are means ± standard error. n=6/experimental group. Main effect of challenge: *p<0.05. Main effect of diet: **p<0.05. Diet/challenge interaction: ***p<0.05. Scheffe post-hoc testing was used for pairwise comparisons (p<0.05).

In male offspring, we observed more differences in transcript abundance in response to CORT challenge (Figure 4A, 4B). In the amygdala, GR decreased in both diet groups (main effect of challenge (F_(3,20)_ = 9.006, p < 0.01), Figure 4C), whereas MR decreased in HFD offspring (main effect of diet (F_(3,20)_ = 4.147, p<0.01), Scheffe post-hoc p=0.047, Figure 4D). Also in the amygdala, with CORT challenge, IκBα (F_(3,20)_ = 52.35, p<0.01, Figure 4E) and IL6 (F_(3,20)_ = 4.82, p = 0.01, Figure. 4F) increased in both CHD and HFD groups. MKP1 increased in HFD offspring (main effect of challenge, (F_(3,20)_ = 9.46, p<0.01), Scheffe post-hoc, p=0.02), but did not change in CHD offspring (Scheffe post-hoc, p=0.170, Figure 4G). In the hippocampus, GR decreased in both diet groups (main effect of challenge, (F_(3,20)_ = 7.69, p<0.01), Figure. 4H), whereas IκBα increased in both diet groups (main effect of challenge, (F_(3,20)_ = 55.3, p<0.01); diet/challenge interaction (F_(3,20)_ = 79.830, p<0.01), Figure 4I). In both diet groups, CORT challenge led to increases in IL6 (F_(3,20)_ = 15.21, p<0.01, Figure 4J) and MKP1 (F_(3,20)_ = 23.41, p<0.01, Figure 4K) in the hippocampus.

**Figure 4.**
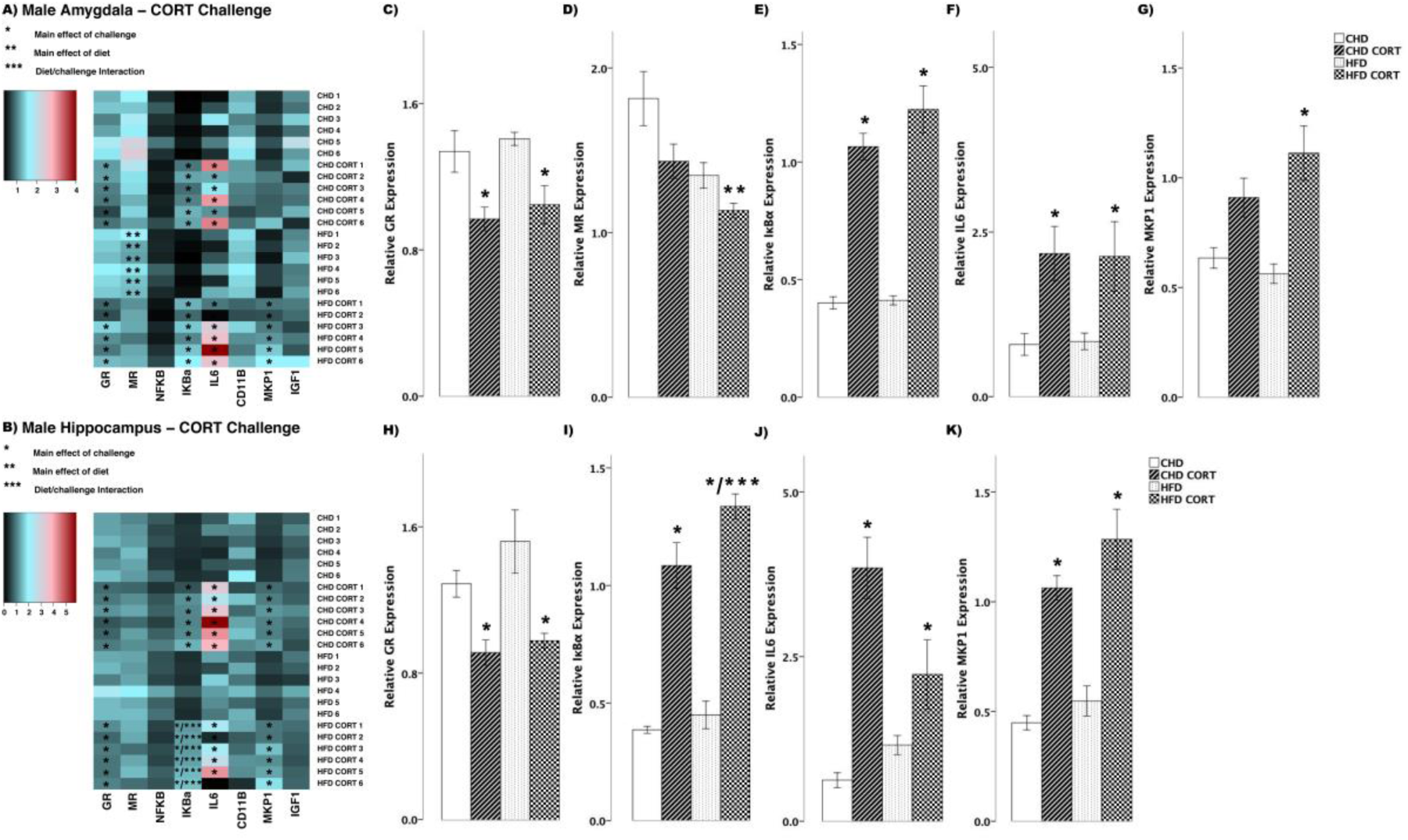
Transcript response to corticosterone (CORT) in the amygdala and hippocampus of adult males. **A and B:** Heatmaps represent transcript levels in the amygdala and hippocampus for individual animals. **C-G:** Relative transcript abundance of GR (glucocorticoid receptor), MR (mineralocorticoid receptor), IκBα (nuclear factor of kappa light polypeptide gene enhancer in B-cells inhibitor, alpha), IL6 (interleukin 6), and MKP1 (mitogen activated protein kinase phosphatase in the amygdala. **H-K:** Relative transcript abundance of GR, IκBα, IL6, and MKP1 in the hippocampus. Data presented are means ± standard error. n=6/experimental group. Main effect of challenge: *p<0.05. Main effect of diet: **p<0.05. Diet/challenge interaction: ***p<0.05. Scheffe post-hoc testing was used for pairwise comparisons (p<0.05).

#### LPS challenge

In female offspring, LPS challenge led to differences in transcript abundance between HFD and CHD offspring (Figure 5A, 5B). Both diet groups showed decreased GR in the amygdala (main effect of challenge (F_(3,20)_ = 6.74, p<0.01), Figure 5C). CHD offspring had decreased MR levels (main effect of challenge F_(3,20)_ = 3.55, p<0.01, Scheffe post-hoc p=0.04), whereas MR levels in HFD offspring remained unchanged (Scheffe post-hoc p=0.749, Figure 5D). Also in the amygdala of both diet groups, there were increases in IκBα (main effect of challenge (F_(3,20)_ = 10.07, p<0.01), Figure 5E), IL6 (F_(3,20)_ = 6.12, p<0.01, Figure 5F), and MKP1 (F_(3,20)_ = 22.63, p<0.01, Figure 5G). In the hippocampus of both diet groups, LPS challenge led to decreased MR transcript (main effect of challenge F_(3,20)_ = 8.70, p<0.01, Figure 5H), while there were increases in NFκB (F_(3,20)_ = 26.80, p<0.01, Figure 5I), and IκBα (F_(3,20)_ = 23.24, p<0.01, Figure 5J). IL6 transcript increased in both diet groups, however, there was a larger increase in HFD females (main effect of challenge (F_(3,20)_ = 6.12, p<0.01); diet/challenge interaction (F_(3,20)_ =2.75, p<0.05, Figure 5K)). MKP1 levels also increased in both diet groups in female offspring (main effect of challenge (F_(3,20)_ = 17.57, p<0.01), Figure 5L). IGF1 transcript levels were lower in HFD females when compared to CHD in basal conditions (main effect of diet (F_(3,20)_ = 2.26, p<0.05), Figure 5M).

**Figure 5.**
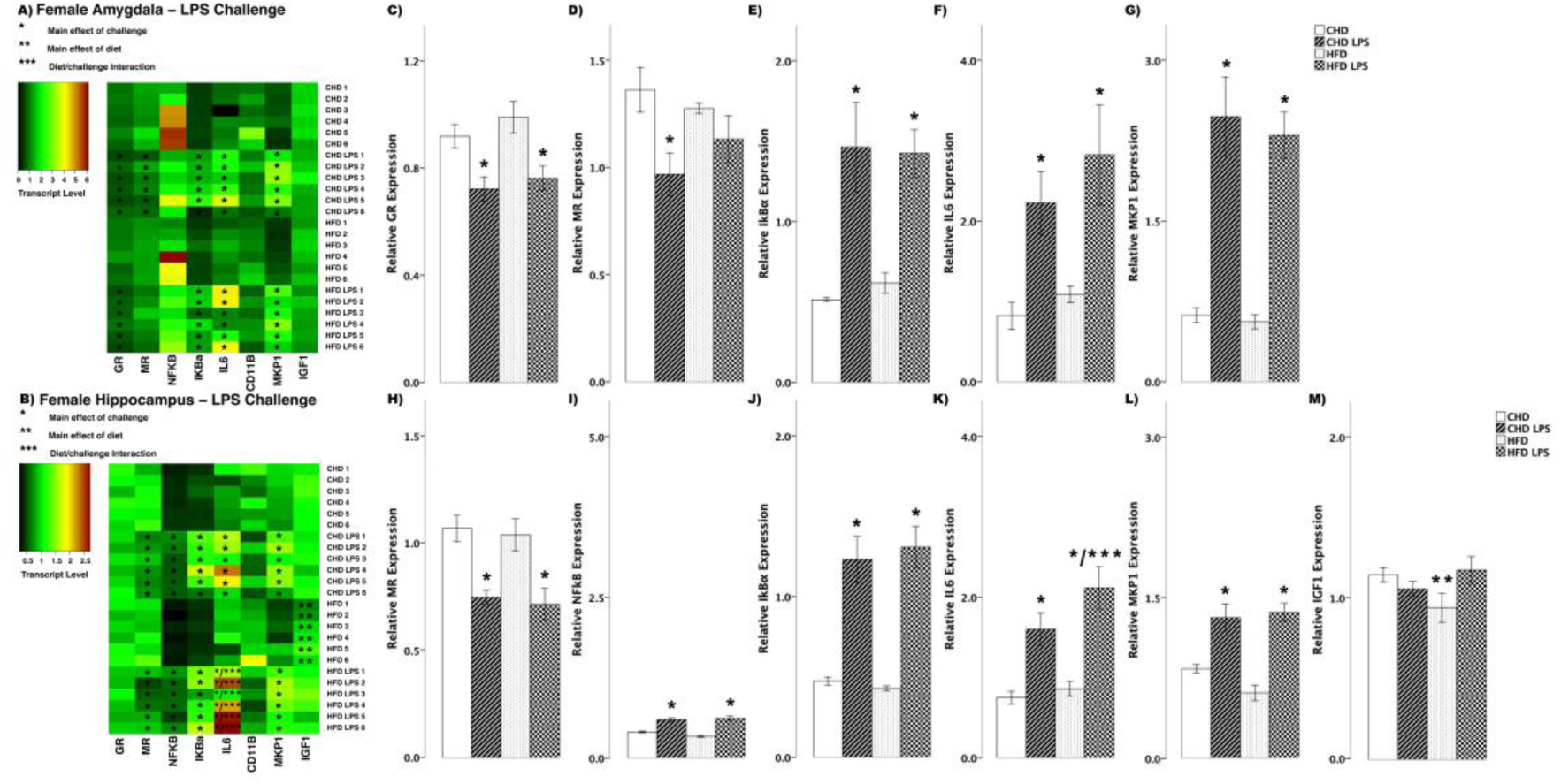
Transcript response to lipopolysaccharide (LPS) in the amygdala and hippocampus of adult females. **A and B:** Heatmaps represent transcript levels in the amygdala and hippocampus for individual animals. **C-G:** Relative transcript abundance of GR (glucocorticoid receptor), MR (mineralocorticoid receptor), IκBα (nuclear factor of kappa light polypeptide gene enhancer in B-cells inhibitor, alpha), IL6 (interleukin 6), and MKP1 (mitogen activated protein kinase phosphatase in the amygdala. **H-M: Relative** transcript abundance of MR, NFκB (nuclear factor kappa light chain enhancer of activated B cells), IκBα, IL6, MKP1, and IGF1 (insulin-like growth factor 1) in the hippocampus. Data presented are means ± standard error. n=6/experimental group. Main effect of challenge: *p<0.05. Main effect of diet: **p<0.05. Diet/challenge interaction: ***p<0.05. Scheffe post-hoc testing was used for pairwise comparisons (p<0.05).

In male offspring, there were fewer differences in transcript abundance between CHD and HFD offspring in response to LPS challenge (Figure 6A, 6B). In the amygdala of both diet groups there were increases in IκBα (main effect of challenge F_(3,20)_ = 28.29, p<0.01, Figure 6C), IL6 (F_(3,20)_ = 7.09, p<0.01, Figure 6D), and MKP1 (F_(3,20)_ = 8.84, p<0.01, Figure 6E) in response to LPS challenge. Similar to changes in the amygdala, in the hippocampus of both diet groups, LPS challenge led to increases in IκBα (main effect of challenge F_(3,20)_ = 18.79, p<0.01, Figure 6F), while IL6 levels decreased in both diet groups (F_(3,20)_ = 5.76, p<0.01, Figure 6G). CD11B (main effect of challenge F_(3,20)_ = 14.95, p<0.01, Figure 6H) and MKP1 (F_(3,20)_ = 9.98, p<0.01, Figure 6I) levels increased in both diet groups in female offspring. Lastly, IGF1 expression decreased in HFD males (main effect of challenge (F_(3,20)_ = 3.73, p<0.05, Scheffe post-hoc p=0.017), but remained unchanged in CHD counterparts (Scheffe post-hoc p=0.410, Figure 6J).

**Figure 6.**
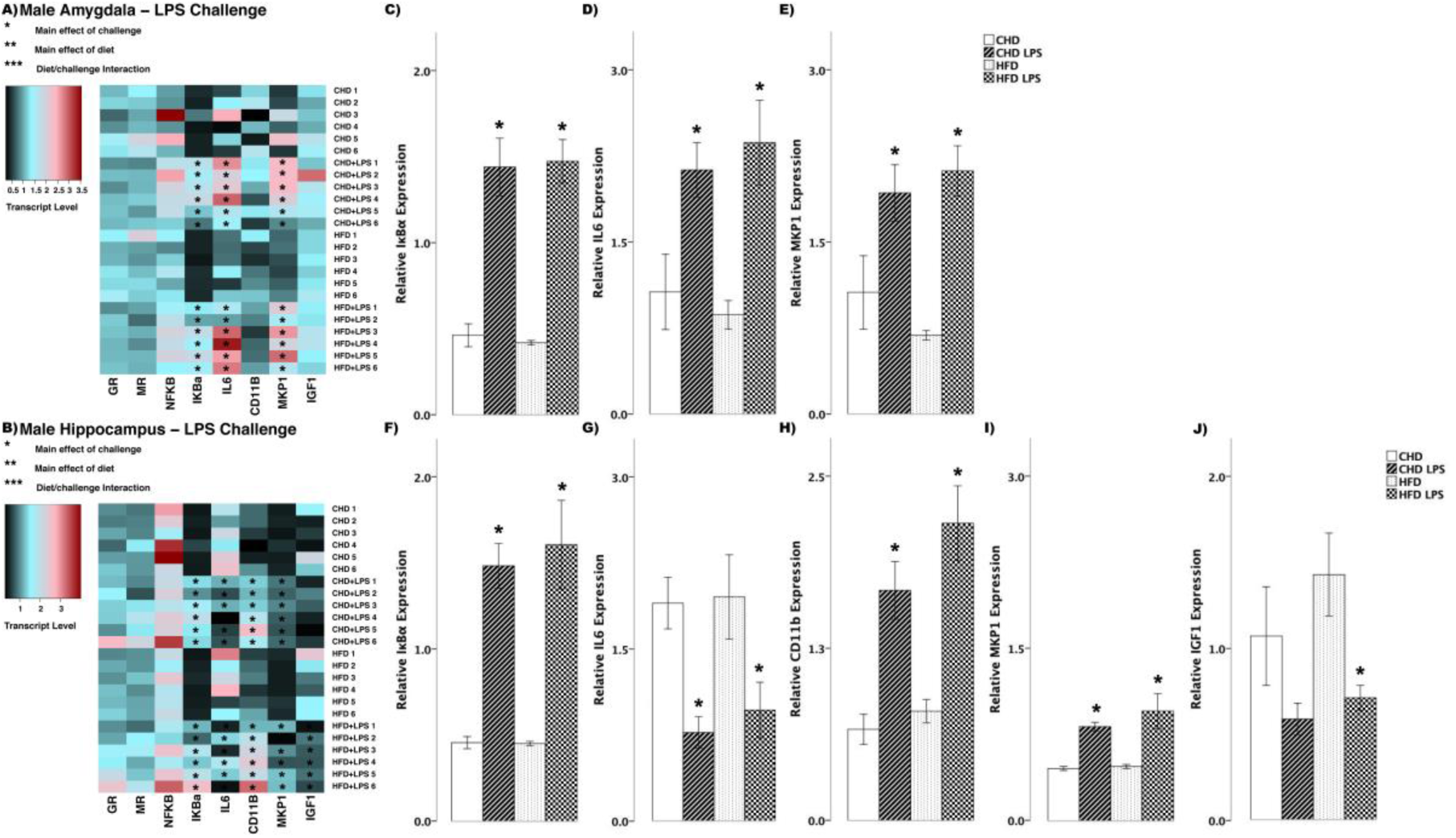
Transcript response to lipopolysaccharide (LPS) in the amygdala and hippocampus of adult males. **A and B:** Heatmaps represent transcript levels in the amygdala and hippocampus for individual animals. **C-E:** Relative transcript abundance of IκBα (nuclear factor of kappa light polypeptide gene enhancer in B-cells inhibitor, alpha), IL6 (interleukin 6), and MKP1 (mitogen activated protein kinase phosphatase 1) in the amygdala. **F-J:** Relative transcript abundance of IκBα, IL6, CD11B (cluster of differentiation molecule 11B), MKP1, and IGF1 (insulin-like growth factor 1) in the hippocampus. Data presented are means ± standard error. n=6/experimental group. Main effect of challenge: *p<0.05. Main effect of diet: **p<0.05. Diet/challenge interaction: ***p<0.05. Scheffe post-hoc testing was used for pairwise comparisons (p<0.05).

#### Combined CORT and LPS challenge

In female offspring, CORT+LPS challenge lead to several differences in transcript abundance between HFD and CHD offspring (Figure 7A and 7B). In the amygdala, both diet groups showed decreases in GR (main effect of challenge (F_(3,20)_ = 10.0, p<0.01), Figure 7C), MR (main effect of challenge (F_(3,20)_ = 9.42, p<0.01); main effect of diet,(F_(3,20)_ = 5.596, p<0.05), Figure 7D), and NFκB (main effect of challenge (F_(3,20)_ = 12.52, p<0.01), Figure 7E). In contrast, IκBα levels increased in HFD offspring (main effect of challenge F_(3,20)_ = 4.68, p<0.05, Scheffe post-hoc p=0.015), but did not change in CHD offspring (Scheffe post-hoc p=0.803, Figure 7F). The ratio of IL6/IL10 was above 1 in both diet groups in the amygdala of female offspring (main effect of challenge (F(3,20) = 9.48, p<0.01); diet/challenge interaction (F_(3,20)=_4.36, p<0.05), Figure 7G). Lastly, CD11B levels decreased in both diet groups (main effect of challenge F_(3,20)_ = 7.19, p<0.01, Figure 7H). In the hippocampus, IκBα (main effect of challenge F_(3,20)_ = 10.13, p<0.01, Figure 7I) and the IL6/IL10 ratio (F_(3,20)_ = 13.06, p<0.01, Figure 7J), increased in both diet groups, while CD11B decreased in CHD females (main effect of challenge F_(3,20)_ = 4.456, p<0.05, Scheffe post-hoc p=0.047) and no change was seen in HFD offspring (Scheffe post-hoc p=0.909, Figure 7K).

**Figure 7.**
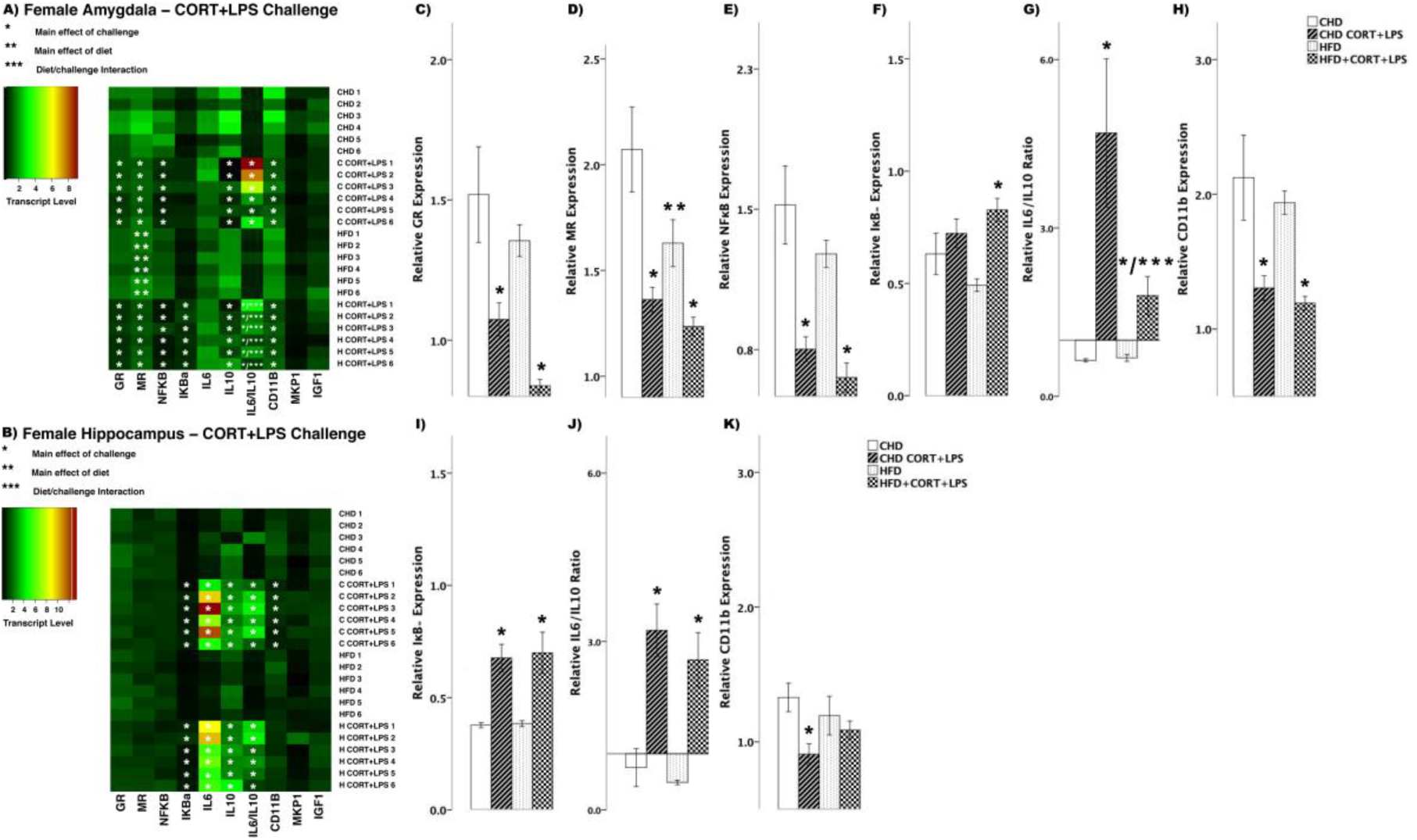
Transcript response to CORT+LPS in the amygdala and hippocampus of adult females. **A and B:** Heatmaps represent transcript levels in the amygdala and hippocampus for individual animals. **C-H:** Relative transcript abundance of GR (glucocorticoid receptor), MR (mineralocorticoid receptor), NFκB (nuclear factor kappa light chain enhancer of activated B cells), IκBα (nuclear factor of kappa light polypeptide gene enhancer in B-cells inhibitor, alpha), the ratio of pro versus anti inflammation designated by IL6/IL10 (interleukin 6/10), and CD11B (cluster of differentiation molecule 11B) expression in the amygdala. **I-K:** Relative transcript abundance of IκB, IL6/IL10 ratio, and CD11B in the hippocampus. Data presented are means ± standard error. n=6/experimental group. Main effect of challenge: *p<0.05. Main effect of diet: **p<0.05. Diet/challenge interaction: ***p<0.05. Scheffe post-hoc testing was used for pairwise comparisons (p<0.05).

In male offspring, there were fewer differences in transcript abundance between diet groups in response to combined CORT+LPS challenge (Figure 8A and 8B). In the amygdala, GR decreased in HFD offspring (main effect of challenge (F_(3,20)_ = 4.364, p<0.05), Scheffe post-hoc p=0.046), yet remained unchanged in CHD offspring (Scheffe post-hoc p=0.512, Figure 8C). MR levels decreased in CHD offspring (main effect of challenge (F_(3,20)_ = 4.110, p<0.05), Scheffe post-hoc p=0.022), yet remained unchanged in HFD offspring (Scheffe post-hoc p=0.356, Figure 8D). NFκB levels decreased in both diet groups (main effect of challenge F_(3,20)_ = 4.44, p<0.05, Figure 8E), where as IL6/IL10 ratio decreased in CHD males (main effect of challenge F_(3,20)_ = 4.458, p<0.05, Scheffe post-hoc p=0.024), but remained unchanged in HFD offspring (Scheffe post-hoc p=0.276, Figure 8F). Lastly, IGF1 levels decreased in both groups in response to CORT+LPS challenge (F_(3,20)_ = 4.29, p<0.05, Figure 8G). In the hippocampus, there was increased GR levels in CHD offspring (main effect of challenge F_(3,20)_ = 4.90, p<0.05, Scheffe post-hoc p=0.044), yet remained unchanged in HFD offspring (Scheffe post-hoc p=0.999, Figure 8H). IκBα transcript levels increased in both diet groups (main effect of challenge F_(3,20)_ = 19.08, p<0.01, Figure 8I). Similar to that of GR, IL6/IL10 ratio increased in CHD offspring (main effect of challenge F_(3,20)_ =5.107, p<0.05, Scheffe post-hoc p=0.025) and did not change in HFD counterparts (Scheffe post-hoc, p=0.319, Figure 8J). Lastly, MKP1 levels increased in both groups (main effect of challenge F_(3,20)_ = 15.42, p<0.01, Figure 8K) in response to CORT+LPS challenge.

**Figure 8.**
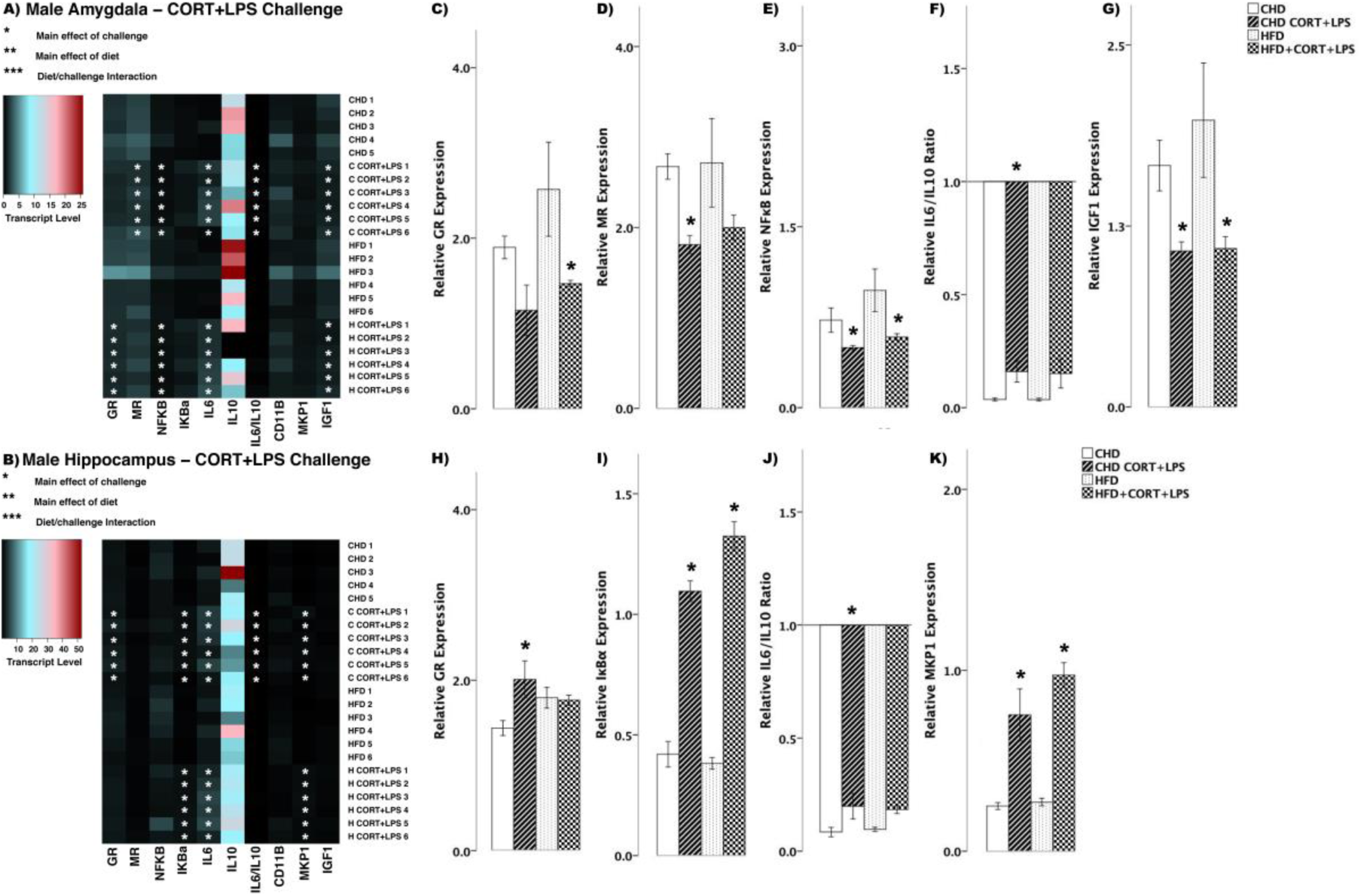
Transcript response to CORT+LPS in the amygdala and hippocampus of adult males. **A and B:** Heatmaps represent transcript levels in the amygdala and hippocampus for individual animals. **C-G:** Relative transcript abundance of GR (glucocorticoid receptor), MR (mineralocorticoid receptor), NFκB (nuclear factor kappa light chain enhancer of activated B cells), the ratio of pro versus anti inflammation designated by IL6/IL10 (interleukin 6/10), and IGF1 expression in the amygdala. **H-K:** Relative transcript abundance of GR, IκBα, IL6/IL10 ratio, and MKP1 in the hippocampus. Data presented are means ± standard error. n=6/experimental group. Main effect of challenge: *p<0.05. Main effect of diet: **p<0.05. Diet/challenge interaction: ***p<0.05. Scheffe post-hoc testing was used for pairwise comparisons (p<0.05).

To better characterize the neural transcript response to combined CORT and LPS exposure, an additional brain region regulating the HPA axis was assessed; the medial prefrontal cortex (PFC; Figure 9A and 9B). In female offspring in response to simultaneous CORT + LPS challenge, both diet groups showed increased levels of NFκB (main effect of challenge F_(3,20)_ = 6.85, p<0.01, Figure. 9C) and IκBα (F_(3,20)_ = 28.95, p<0.01, Figure 9D), while IL6 increased in HFD offspring (main effect of challenge F_(3,20)_ = 4.347, p<0.05, Scheffe post-hoc p=0.040), but did not change in CHD offspring (Scheffe post-hoc p=0.784, Figure 9E). MKP1 levels increased in HFD offspring with CORT+LPS challenge (diet/challenge interaction (F_(3,20)_ = 2.08, p<0.05), Figure 9F) when compared to CHD counterparts.

**Figure 9.**
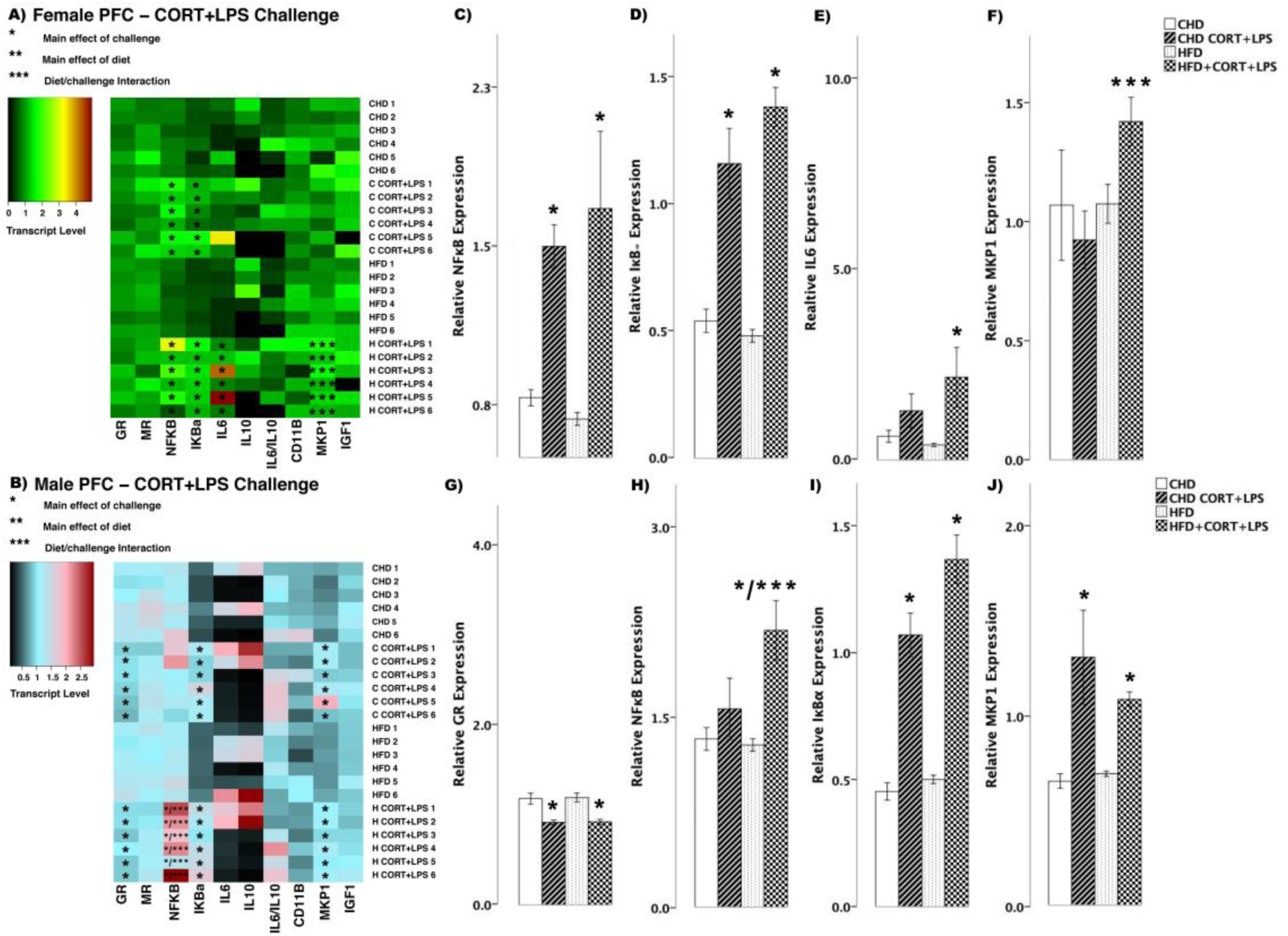
Transcript response to CORT+LPS in the prefrontal cortex of adult females and males. **A and B:** Heatmaps represent transcript levels in the PFC for individual female and male animals. **C-F:** Relative transcript abundance of NFκB (nuclear factor kappa light chain enhancer of activated B cells), IκBα (nuclear factor of kappa light polypeptide gene enhancer in B-cells inhibitor, alpha), IL6 (interleukin 6), and MKP1 (mitogen activated protein kinase phosphatase 1) in the PFC of females. **G-J:** Relative transcript abundance of GR, NFκB, IκBα, and MKP1 in the PFC of males. Data presented are means ± standard error. n=6/experimental group. Main effect of challenge: *p<0.05. Main effect of diet: **p<0.05. Diet/challenge interaction: ***p<0.05. Scheffe post-hoc testing was used for pairwise comparisons (p<0.05).

In male offspring, GR levels decreased in the PFC of both diet groups (F_(3,20)_ = 13.45, p<0.01, Figure 9G) in response to combined CORT and LPS treatment. NFκB levels increased in HFD males (main effect of challenge (F_(3,20)_ = 8.111, p<0.01, Scheffe post-hoc p=0.007); diet/challenge interaction (F_(3,20)_ = 5.486, p<0.01), whereas no change was seen in CHD offspring (Scheffe post-hoc p=0.994, Figure 9H). Lastly, CORT+LPS challenge led to increased IκBα (F_(3,20)_ = 42.56, p<0.01, Figure 9I) and MKP1 (F_(3,20)_ = 9.53, p<0.01, Figure 9J) levels in both CHD and HFD male offspring.

## DISCUSSION

Overall, and as expected based upon previous findings, CORT potentiated pro-inflammatory gene expression induced by LPS in both female and male CHD offspring. This potentiation was also observed in HFD offspring, however compared to the CHD animals, there were elevated levels of pro-inflammatory transcript in males, while females exhibited elevations in anti-inflammatory transcript (Figure 10). These findings indicate distinct transcriptional responses to elevated CORT and LPS among HFD-exposed offspring, some of which were sex-specific.

**Figure 10.**
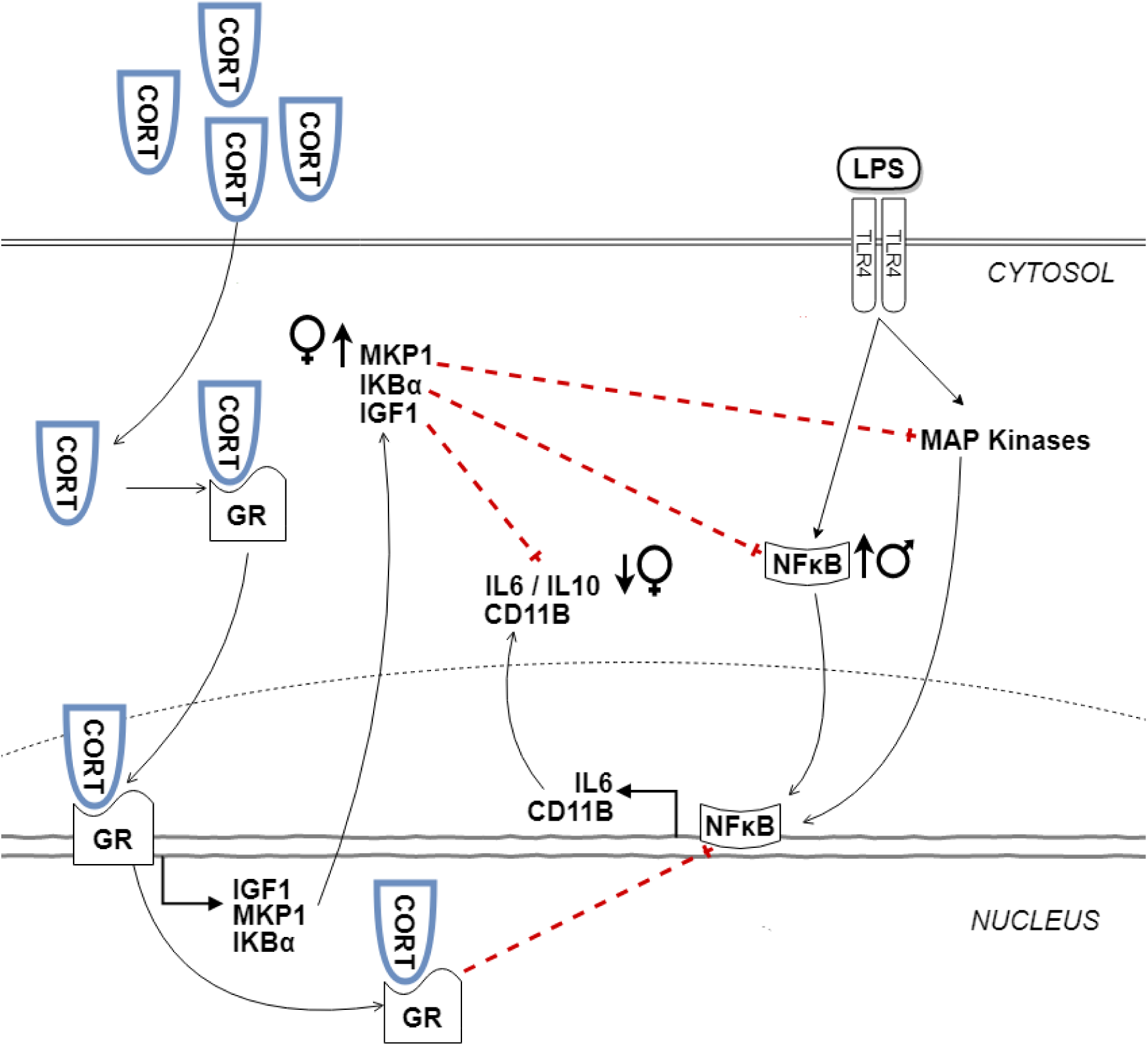
Maternal high-fat diet induces sex-specific effects on transcript responses to CORT+LPS challenge in adulthood. CORT diffuses into the cytosol and binds to glucocorticoid receptor (GR). The CORT-GR complex induces the expression of anti-inflammatory molecules including IGF1, MKP1, and IKBα, which at basal levels inhibit pro-inflammatory expression of NFκB, IL6 and CD11B induced by LPS signalling to toll-like receptor 4 (TLR4) and mitogen-activated protein (MAP) kinases. However, with higher levels of CORT, anti-inflammatory effects are reduced. This system was altered in females exposed to maternal high-fat diet (HFD), as indicated by was increased MKP1 and a reduced IL6/IL10 ratio. Males exposed to maternal HFD showed increased NFκB transcript.

### Enhanced anti-inflammatory responses to CORT challenge in male HFD offspring

CORT is classically associated with anti-inflammatory and immunosuppressive effects. The anti-inflammatory actions of CORT include increased expression of IGF-1, IκBα, and MKP1, which supress the expression of inflammatory mediators (34,35). In this study, both diet groups and sex showed increased expression of IκBα in the hippocampus in response to CORT challenge (Figures 3-4). However, in males, the increase in IκBα in the hippocampus in response to CORT was greater among HFD offspring when compared to CHD offspring. Overall, HFD males also showed reduced MR transcript in the hippocampus. It is possible that reduced MR transcript may have potentiated further GR-induced anti-inflammation with CORT administration, since reduced availability of MR and receptor saturation with CORT is associated with increased GR expression (35–37). This would suggest that male offspring exposed to maternal HFD may tolerate acute psychological or endocrine stressors more efficiently than CHD offspring in adulthood, initiating protective, anti-inflammatory signalling in the hippocampus.

### Enhanced pro-inflammatory response to LPS challenge in female HFD offspring

LPS binds to toll-like receptor 4, leading to the activation of MAP kinases that support inflammatory gene transcription. Over time, pro-inflammatory activation of the HPA leads to a homeostatic state through GR-mediated expression of the anti-inflammatory genes MKP-1 and IκBα that prevent further accumulation of cytotoxic pro-inflammatory cytokines (1,38–42). We found that LPS challenge largely led to increases in both pro- and anti-inflammatory transcripts in both sexes and diet groups, with a potentiated IL6 response in the hippocampus of HFD females (Figures 5-6). Across sexes, diet groups, and brain regions, we found increased anti-inflammatory IκBα and MKP1 in response to LPS. Both sexes and diet groups also showed increased IL6 transcript in the amygdala, and females showed increased NFκB transcript in the hippocampus in response to LPS. In the hippocampus however, we found that females exposed to maternal HFD had significantly higher levels of IL6 than CHD controls post-LPS. The distinct IL6 response in the hippocampus of female HFD offspring indicates that maternal HFD exposure potentiates inflammatory mechanisms in HFD offspring in conditions of immune stress. Notably, similar results have been reported in the hypothalamus, where chronic HFD consumption leads to increased inflammation and neuronal apoptosis (43,44).

### Sex-specific alterations to CORT+LPS challenge in HFD offspring

Basal levels of CORT are known to dampen the pro-inflammatory effects of LPS through GC signalling and anti-inflammatory gene expression. A previous study reported acute elevations in CORT through exogenous administration or psychosocial stress increases pro-inflammatory transcript expression, but decreases anti-inflammatory transcript expression (3,5). In the amygdala of female offspring exposed to maternal HFD, we found that simultaneous CORT and LPS exposure led to a lower pro-/anti-inflammatory IL6/IL10 cytokine transcript ratio relative to CHD females, indicating an enhanced anti-inflammatory transcriptional response (Figure 7). In contrast, simultaneous CORT and LPS exposure in males led to minimal differences in the amygdala and hippocampus between the diet groups (Figure 8). In the PFC, only HFD females displayed an increase in MKP-1 (Figure 9). These findings suggest that maternal HFD exposure is predominantly associated with increased anti-inflammatory transcriptional response in the amygdala and PFC of females. In contrast, CORT and LPS exposure led to a significant increase in NFκB in the PFC only in HFD males (Figure 9). These findings suggest that maternal exposure to HFD exacerbates neuroinflammatory transcriptional responses to combined CORT and LPS challenge in the PFC of male adults.

It is tempting to speculate that these enhanced pro-inflammatory transcript responses may arise as an adaptive response to perinatal stress induced by maternal HFD exposure (45). In the context of maternal obesity induced by high levels of saturated fat, there are elevations in circulating fat and adiposity that stimulate inflammation, which in turn stimulates CORT elevations through HPA axis activation (10,11). In the mother, elevated CORT and inflammatory molecules can be transmitted to developing offspring during fetal and lactation stages (46–48). It is possible that exposures to elevated levels of CORT and inflammation during these critical points in development program HPA axis activity and glucocorticoid signalling in offspring confer adaptation to chronic inflammation experienced in perinatal life (19–25). Inasmuch as these alterations based on perinatal cues of inflammation do not match the environment in post-natal life, they may constitute a mechanism leading to phenotypic changes in behavior. Consistent with this hypothesis, elevated anxiety behavior is associated with pro-inflammatory gene expression and inflammation in brain regions regulating the HPA axis, such as the amygdala, hippocampus, and PFC (49,50). In this study, the hippocampus was the site of enhanced anti-inflammatory responses post-CORT in HFD males and pronounced pro-inflammatory responses post-LPS in HFD females. With CORT+LPS challenge, males showed enhanced pro-inflammatory responses in the PFC, while females showed the opposite in both the PFC and amygdala. Future studies are necessary to assess how the sex- and brain region-specific alterations to the inflammatory transcript response to CORT andLPS observed herein alter anxiety-like behavior in offspring with perinatal HFD exposure.

## CONCLUSION

In this study, we found that maternal HFD exposure altered transcript responses in adult offspring, through enhanced anti-inflammation post-CORT challenge in males and enhanced pro-inflammation post-LPS in females. We also found evidence of enhanced pro-inflammation post-CORT+LPS challenge in males, with females exhibiting enhanced anti-inflammation. Our findings suggest that exposure to maternal HFD during early life induces sex-specific transcriptional responses to stress in adulthood. It is currently unknown whether humans exposed to maternal obesity and high-fat diets may show similar responses to glucocorticoid and immune stress. However, past studies in rodents and humans indicating that obesity leads to prolonged inflammation and increased sensitivity to infection, suggesting that these effects should also be examined in the context of humans exposed to obesity during development (51–53).

## AVAILABILITY OF DATA AND MATERIALS

The data used in this study are available from the corresponding author upon request.

## ABBREVIATIONS

ANOVA: Analysis of variance
CD11B: Cluster of differentiation molecule B
CHD: Control house-chow diet
CORT: Corticosterone
GAPDH: Glyceraldehyde 3-phosphate dehydrogenase
HFD: High-fat diet
HPA: Hypothalamic-pituitary-adrenal
HPC: Hippocampus
IGF1: Insulin-like growth factor 1
IκBα: Nuclear factor of kappa light polypeptide gene enhancer in B-cells inhibitor, alpha
IL6: Interleukin 6
IL10: Interleukin 10
GR: Glucocorticoid receptor
LPS: Lipopolysaccharide
MKP1: Mitogen-activated protein kinase phosphatase 1
MR: Mineralocorticoid receptor
NFκB: Nuclear factor kappa-beta-light-chain-enhancer of activated B cells
PFC: Prefrontal cortex
PND: Post-natal day
qPCR: Quantitative polymerase chain reaction
SEM: Standard error of mean
YWHAZ: Tyrosine 3-monooxygenase/tryptophan 5-monooxygenase Activation Protein Zeta

## ACKNOWLEDGEMENTS

We thank Dr. Sameera Abuaish and Shathveekan Sivanathan for helping with molecular and animal work, and Dr. Sohee Kang, Associate Professor and statistics coordinator at the University of Toronto Scarborough, for providing statistical advice and guidance.

## AUTHORS’ CONTRIBUTIONS

AS and POM designed the experiment. AS, POM, and WCDV conducted the animal work. CMWL, WCDV and MFR conducted the molecular work. SW and MFR analyzed the data. MFR, SW and POM interpreted the data and wrote the manuscript. All authors read and approved the final manuscript.

## ETHICS DECLARATION

### Ethics approval and consent to participate

All animal studies complied with the Canadian Council on Animal Care Guidelines and Policies and were approved by the Local Animal Care Committee at the University of Toronto Scarborough.

### Consent for publication

Not applicable.

